# Hippocampal ripples coincide with “up-state” and cortical spindles in Retrosplenial Cortex

**DOI:** 10.1101/2022.12.19.521088

**Authors:** Rafael Pedrosa, Mojtaba Nazari, Loig Kergoat, Christophe Bernard, Majid Mohajerani, Federico Stella, Francesco Battaglia

**Author notes:** **Correspondence:** Rafael Pedrosa, Ms.C., Francesco Battaglia, Ph.D., Radboud University, Donders Institute for Brain, Cognition and Behaviour, Neuroinformatica, Heyendaalseweg 135, 6525 AJ NIJMEGEN, Internal postal code: 66, and.

## Abstract

During NREM sleep hippocampal Sharp-wave ripples (SWR) events are thought to stabilize memory traces for long-term storage in downstream neocortical structures. Within the neocortex, Default Mode Network (DMN) areas interact preferentially with the hippocampus purportedly to consolidate those traces. Transient bouts of slow oscillations and sleep spindles in DMN areas are often observed around SWRs, suggesting that these two activities are related and that their interplay possibly contributes to memory consolidation. To investigate how SWRs interact with the DMN and spindles, we combined cortical wide-field voltage imaging, ECoG, and hippocampal LFP recordings in anesthetized and sleeping mice. Here we show that, during SWR, “up-states” and spindles reliably co-occur in a cortical subnetwork centered around the Retrosplenial cortex. Furthermore, Retrosplenial transient activations and spindles predict Slow Gamma oscillations in CA1 during SWRs. Together, our results suggest that Retrosplenial-hippocampal interaction may be a central source of information exchange between cortex and hippocampus.

## Introduction

The consolidation of newly encoded memories, facilitating their long-term retention, can occur during sleep (Born and Wilhelm, 2012; Stickgold and Walker, 2005), in particular though an hippocampal-cortical dialog during Non-rapid eye movement sleep (NREM)(Girardeau et al., 2014; Rolls, 2000; Wang and Morris, 2010). The replay of neural ensembles in the hippocampus (Foster and Wilson, 2006; Nádasdy et al., 1999; Ólafsdóttir et al., 2016; Wilson and McNaughton, 1993), as well as the repeated activation of cortical patterns (Higgins et al., 2021; Peigneux et al., 2004; Tambini and Davachi, 2019) support this view. However, the physiological and dynamical properties of this interaction remain largely unknown. Several arguments support the involvement of Sharp-Wave Ripples (SWR, 100-200 Hz), a property of the CA3-CA1 network (Buzsáki, 2015), during sleep: 1) Support the replay of neural ensembles that have been engaged in memory encoding (Foster and Wilson, 2006; Peyrache et al., 2009), 2) the suppression sleep SWR impairs memory performance (Girardeau et al., 2009; Maingret et al., 2016), and 3) SWRs are coupled to the active, UP-state of Slow Oscillation (SO; <1Hz) in specific associative areas in the neocortex (Battaglia et al., 2004; Sirota et al., 2003).

Another ‘spontaneous’ oscillatory component observed on the cortex during NREM periods are sleep spindles (Ngo et al., 2020; Siapas and Wilson, 1998). These events are waxing and waning oscillatory (8-16 Hz) bouts generated in the thalamus, triggered by the depolarizing inputs generated by cortical UP states (Timofeev and Steriade, 1996). These events are also thought to be a component of memory consolidation across different species (Cowan et al., 2020; Johnson et al., 2010; Latchoumane et al., 2017; Xu et al., 2021), and they co-occur with the hippocampal SWR (Ngo et al., 2020; Siapas and Wilson, 1998), as well as the UP state of the SO on the cortical surface (Niethard et al., 2018). Taken together, the SO-spindle-SWR loop induce synchronized neural plasticity, which may participate in the mechanisms of memory consolidation (Niethard et al., 2018; Staresina et al., 2015).

The Default Mode Network (DMN) is a resting-state network, which among other functions, has been implicated in memory processes (Buckner et al., 2008; Tambini et al., 2010). Recently, combining electrophysiology in the hippocampus with brain Blood Oxygen Level Dependent (BOLD) activity, Kaplan et al. reported that primate SWRs were related to a strongly correlated pattern of activity over regions of the DMN (Kaplan et al., 2016). More recently, this result has also been confirmed in mice on a finer time scale with wide-field voltage imaging (Karimi Abadchi et al., 2020; Pedrosa et al., 2022a). In rodents, the DMN comprises areas such as: the Orbitofrontal, Anterior Cingulate, Retrosplenial, Parietal, Temporal Association Cortex and in addition the dorsal hippocampus (CA1, (Lu et al., 2012). Studies investigating the functional connectivity between SWRs and these DMN regions, has shown a diversity of spatiotemporal patterns, which implies that SWR may interact with cortex in multiple modes (Karimi Abadchi et al., 2020; Oliva et al., 2016; Opalka et al., 2020; Wang and Ikemoto, 2016).

In fact, little is known about the relationship between spindles in the DMN and hippocampal SWRs, a question that requires better spatiotemporal resolution than fMRI to be addressed. In this study, we combined wide-field Voltage Sensitive Dye (VSD) imaging and Electrocorticogram (ECoG) from the neocortex with Local Field Potential (LFP) recording from the ipsilateral CA1 hippocampus. We discovered different cortical patterns of SO within the DMN and across the dorsal cortex and we studied them in the context of their interaction with hippocampal SWRs during NREM sleep. Interestingly, we found that a pattern centered in Retrosplenial cortex which is topographically matched by the spatial distribution of cortical Spindles. In addition, we found that both SO and spindle transients in Retrosplenial cortex are correlated with Slow Gamma (20-45 Hz) in CA1 *stratum radiatum*. Combined SO-Spindle transients observed in Retrosplenial Cortex tend to precede hippocampal SWR, suggesting that Retrosplenial cortex is a hub of spontaneous activity processes during sleep, potentially with a key role for memory consolidation.

## Results

### Experimental design for the spatiotemporal investigation of the neocortex during SWR event

To investigate the neocortical activity pattern during hippocampal SWRs, we used a similar experimental protocol developed previously by Mohajerani and colleagues (Karimi Abadchi et al., 2020), combining Voltage Sensitive Dye (VSD) imaging (Bermudez-Contreras et al., 2018; Kyweriga et al., 2017; Mohajerani et al., 2013) and electrophysiology in the dorsal hippocampus of mice under urethane anesthesia (Figure 1 A-D). The location of primary visual and select somatosensory areas was determined with the help of sensory stimulation and the location in the image of all cortical areas was estimated relative to those areas (Figure 1 B; See methods)(Mohajerani et al., 2010).

**Figure 1:**
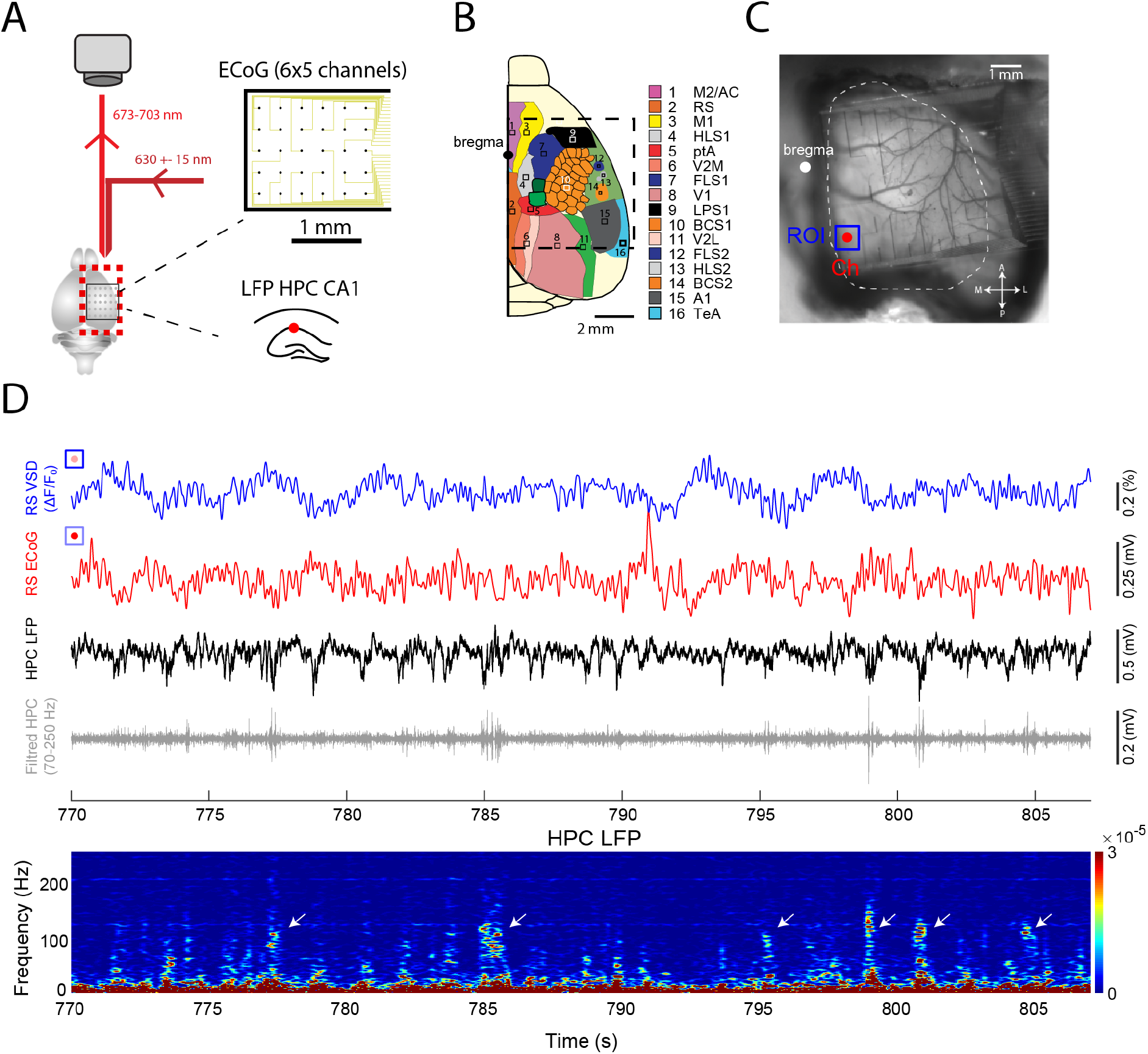
Experimental setup used for recording the neocortical activity around SWRs. **(A)** Schematic of the experimental setup used for combined neocortical optical imaging, hippocampal LFP and electrocorticography. The VSD imaging was performed using a CCD camera Ipsilateral LFP of the dorsal CA1 was recorded from a single electrode implanted from the posterior region of the scalp. A neocortical LFP was also recorded from a transparent grid of electrodes (6×5 channels). **(B)** Illustrative neocortical topography for wide-field optical imaging. The black dashed line represents the boundary of the imaging window. Bregma is represented as a black dot. **(C)** Photograph of a wide unilateral craniotomy with the transparent ECoG grid. The white dashed contour renders the extent of the neocortex. Bregma is depicted as a white dot. Axes: anterior (A), posterior (P), medial (M) and lateral (L). The red dot is an ECoG recording site on the Retrosplenial cortex and the blue square is a Region of Interest (ROI) selected around this Channel (Ch). **(D)** Example of the optical (blue) and ECoG (red) signals from the Retrosplenial cortex selected from the ROI and recording site shown in (C). Note that both activities have a similar oscillatory pattern. Both VSD and ECoG signals are filtered between 0.2 and 6 Hz. Below, aligned hippocampal LFP (black) and filtered hippocampal LFP (gray, 70-250 Hz). Bottom, the spectrogram of the hippocampal LFP shows peaks of high frequency oscillation.

In addition, we recorded local electrocorticograms (Figure S1 A-B) (ECoG; 6×5 matrix with a pitch of 1 mm between the electrodes) from an electrode grid based on the conformable and transparent parylene C substrate and the mixed organic ionic/electronic conductor, poly(3,4-ethylenedioxythiophene):polystyrene sulfonate (PEDOT:PSS). PEDOT:PSS is commonly used to lower impedance of metallic electrodes, typically by one order of magnitude (Sessolo et al., 2013), leading to higher signal-to-noise ratio (SNR). The impedance of our electrodes was about 20 kΩ for electrodes of 20 μm x 20 μm (Figure S1 C). PEDOT:PSS-based ECOGs have been used on top of the brain to record LFPs and even action potentials in rodents and humans (Khodagholy et al., 2016, 2015, 2013). Higher SNR also means that we can use smaller electrodes, which, combined with the transparency of the substrate, leads to a better optical imaging due to less shadow from the electrodes in the field of view (Donahue et al., 2018). The ECOG was designed to cover the entire cortical area recorded (Figure 1 C).

### Hippocampal SWRs in mice is associated with Default Mode Network activity in different temporal scales

First, we investigated the spatiotemporal activity around the detected SWR events (150-300 Hz) in the hippocampus CA1. As previously reported, at the time of the SWR appearance, the cortical areas involved in the DMN became active (Figure 2 A; Video 1)(Kaplan et al., 2016; Karimi Abadchi et al., 2020). In particular, areas along the midline activated, namely Retrosplenial (RS) and Anterior Cingulate cortices (AC), together with a lateral complex of areas involving the Primary Auditory cortex with the Temporal Association cortex (A1/TeA) (Figure 2 B). Here, because the proximity between AC and Secondary Motor cortex makes it difficult to resolve these two areas in our technique, we referred to this area as M2/AC (Karimi Abadchi et al., 2020; Mohajerani et al., 2013). This result could also be observed across animals (Figure 2 C).

**Figure 2:**
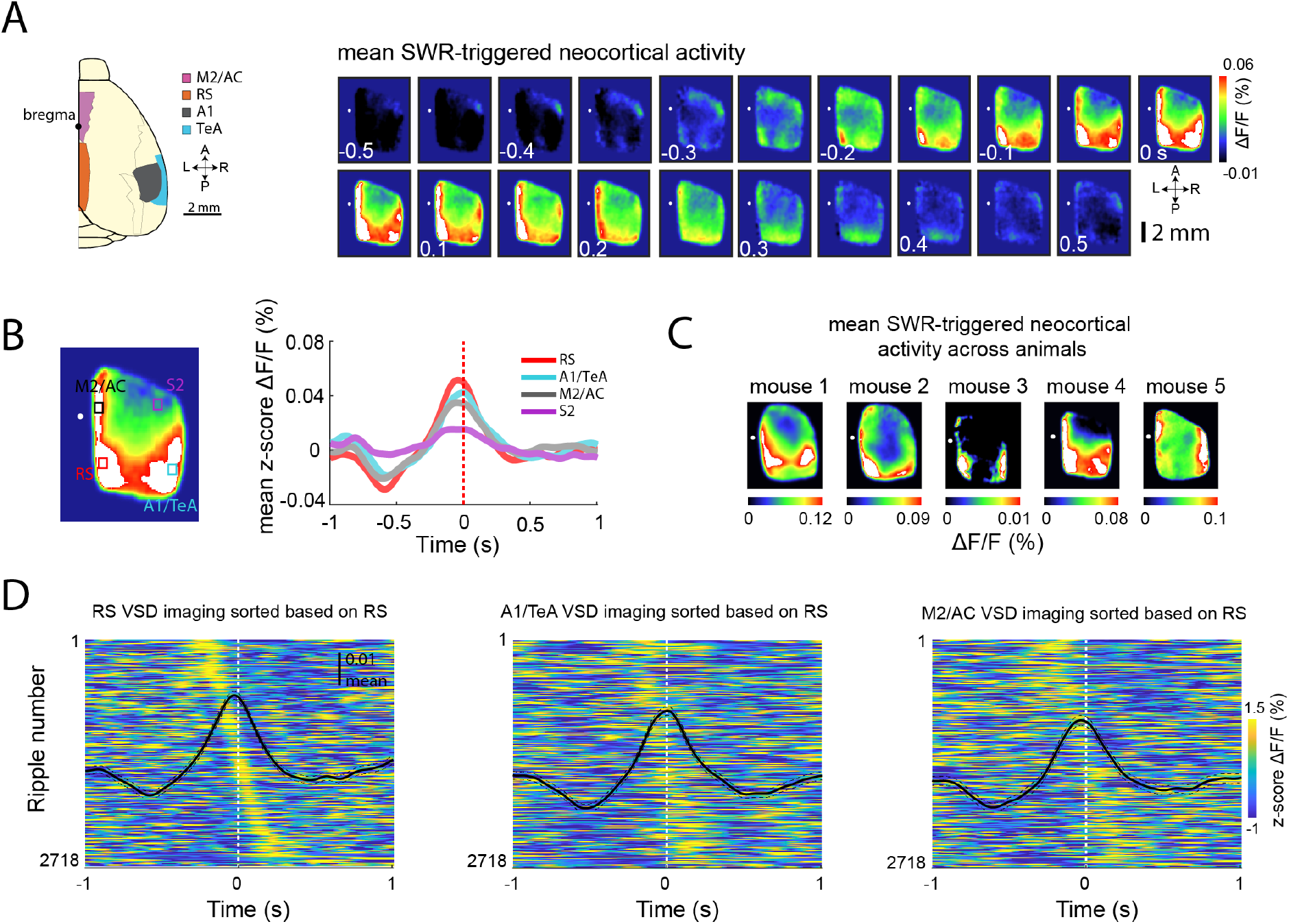
DMN areas activate differentially around hippocampal SWR. **(A)** Mean cortical voltage imaging signal around SWR events. The white dot represents the bregma position. **(B)** On left, the averaged cortical activity around SWR (between −100 ms and 100 ms) from the example in (A). The selected areas are: Retrosplenial (RS, in red), Secondary Motor and Anterior Cingulate (M2/AC, in black), Secondary Sensory (S2 in purple) and Primary Auditory and Temporal Association Area (A1/TeA, in cyan). For each mouse we determined the ROI based on the evoked activity area from the stimulation data and the topographic map (see Material and Methods). On the right, the averaged activity across animals of areas shown on the left panel (n=5 mice). **(C)** Averaged cortical activity (between −100 ms and 100 ms) for each individual mouse. Note for all the mice the high activity of the DMN areas (mouse 4 is the animal from the example in (A)). **(D)** Aligned RS, A1/TeA and M2/AC activity around all the detected SWR event. The plots are sorted by the RS activity.

Although peak activation of RS, A1/TeA and M2/AC around the hippocampal SWRs are significantly correlated (Figure 2 D; Total: 2718 SWRs detected. p<0.0001 for all comparisons, using Spearman’s Rank correlation), there is considerable variability across events and areas, suggesting that DMN responses to SWRs may be more diverse than previously reported (Karimi Abadchi et al., 2020; Pedrosa et al., 2022a).

### Multi-modal Default Mode Network activation around SWRs

To investigate the dynamic patterns of the cortical DMN areas around the hippocampal SWRs, we used unsupervised classification to find the canonical cortical activation modes that accompanied the hippocampal SWR event. First, we applied Register Multimodal Images (RMI) on the cortical activity (averaged from −0.2 to 0.2 s) for all the detected SWR to apply a common mask (Figure 3 A) and merge data from multiple animals. Next, we applied principal component analysis (PCA) followed by independent component analysis (ICA) to these cortical activity maps SWRs (Figure 3 B). As result, we observed a separation of the DMN areas across components (Figure 3 C), which was independent of the number of extracted ICA components (Figure S2).

**Figure 3:**
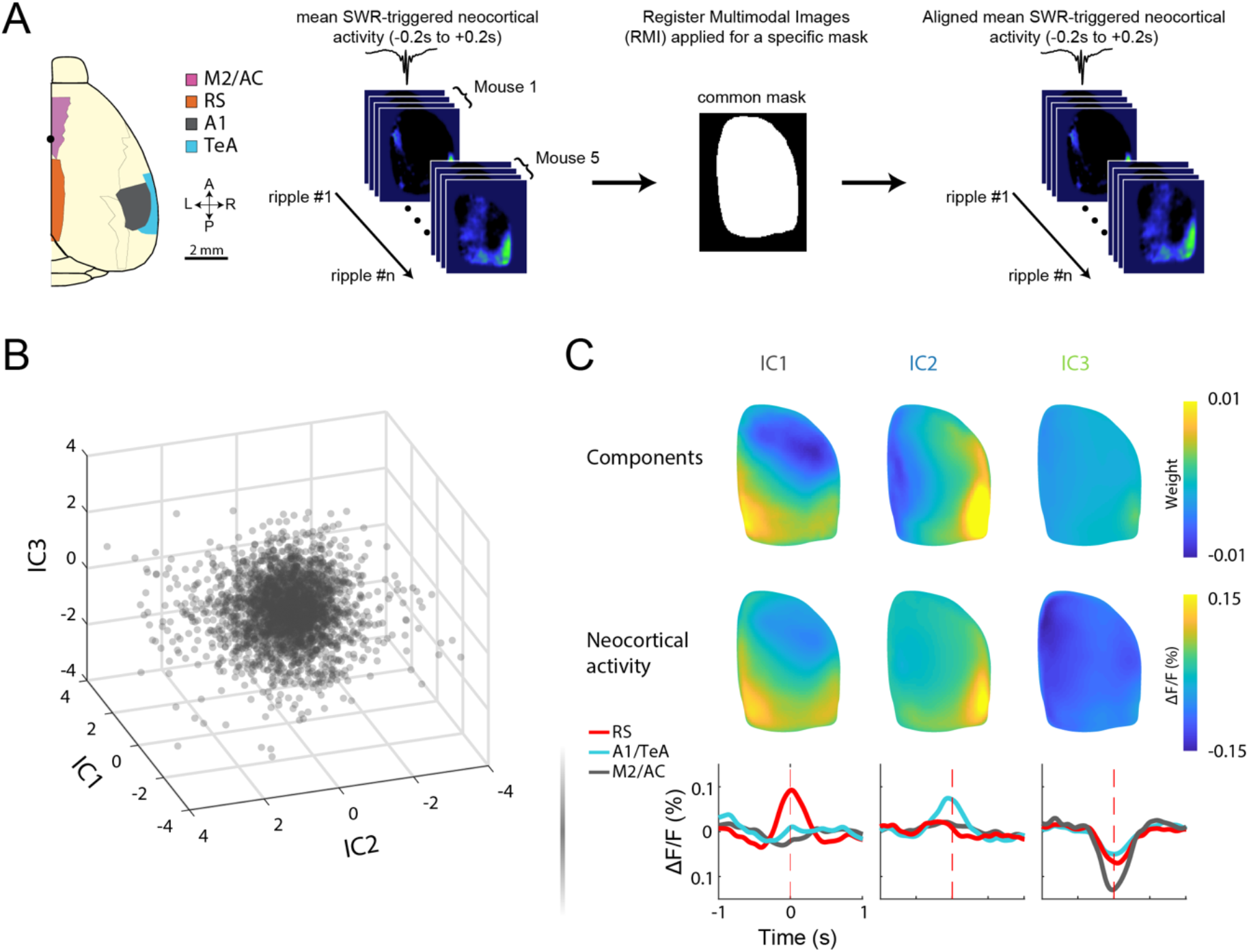
DMN has distinct cortical activity patterns around hippocampal SWR. **(A)** Schematics of the pipeline used to merge data across animals. For the cortical map characterization, we averaged the cortical imaging data between −0.2 to 0.2 s around each detected SWR **(B).** Scatter plot shows the first 3 estimated independent components of the cortical activity for each detected SWR (n=2718 from 5 mice). **(C)** ICA components (top) and the averaged neocortical activity from the 20% of the events with highest weight for each IC (middle). On the bottom, we show the time evolution of RS, A1/TeA and M2/AC from the averaged events showed in the middle plot.

Strikingly, IC1 and IC2 showed a partition of the DMN as reported in Figure 2, between the activity of, respectively, RS and M2/AC from A1/TeA. So far, only patterns compatible with IC1 has been previously reported in the literature during sleep condition (Karimi Abadchi et al., 2020; Pedrosa et al., 2022a). IC3 showed instead an inhibitory pattern across multiple cortical areas. We next investigated how the properties of SWR events varied according to the corresponding values of the cortical ICA scores. Interestingly, we found a significant correlation between score of the ICA vs Ripple amplitude for IC2 and IC3, while for IC1 we found a significant correlation with the Ripple duration (n=2718, *p<0.05; Figure S2 B). We did not find significant correlation with ripple frequency. Together, these findings indicate that features of SWR events might correlate with DMN activity.

Although the number of SWRs varies across mouse (Mouse 1 = 177, Mouse 2 = 669, Mouse 3 = 1000, Mouse 4 = 423 and Mouse 5 = 449. Total = 2718), the distribution of events dominated by each cortical IC for each animal is similar (Figure S2 C-D). To measure this event distribution, we distinguished 3 groups of cortical activity by clustering the vector of per-event IC scores with k-means using data from all mice (Figure S2 C; distribution of events per group: Group1 = 42%, Group2 = 26% and Group3 = 32%). The events classified in each group (equivalent to the IC’s from Figure 3 C) were then reassigned to the respective mouse (Figure S2 D).

### Sleep spindles are mostly activated in RS around SWRs

Although, as shown above and previously in the literature (Battaglia et al., 2004; Siapas and Wilson, 1998) SWR, sleep spindle and slow oscillations have a tendency to co-occur, little is known the spatiotemporal features of their interaction. To characterize this relation, we used the ECoG data to analyze the sleep spindle activity around the SWR during the different cortical IC groups (Figure 4 A).

**Figure 4:**
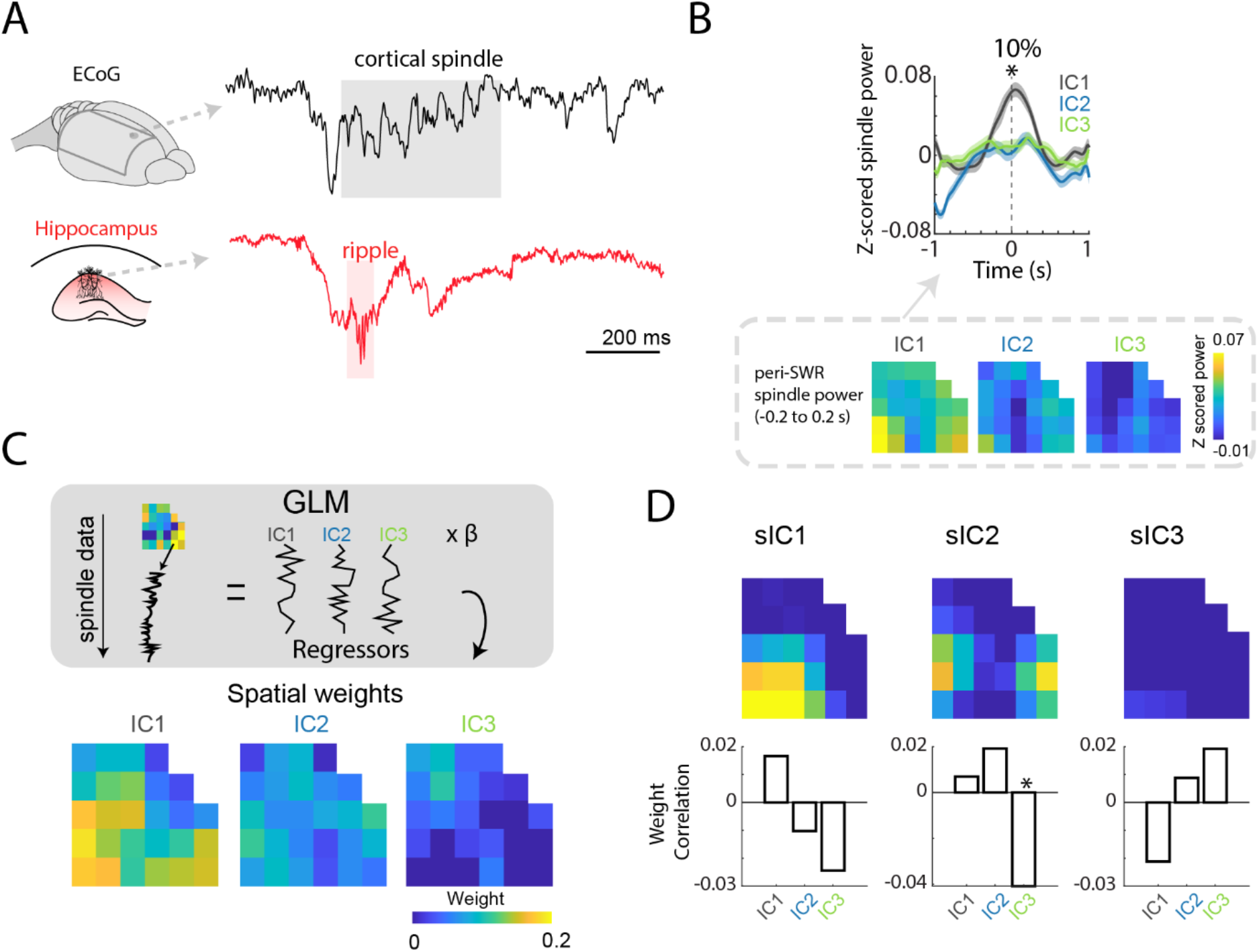
Cortical spindle activity coactivate with SO in RS around hippocampal SWR. **(A)** Example of a SWR event detected in the hippocampus coupled with a cortical spindle detected in a frontal electrode from the ECoG. **(B)** On top, the global spindle power (8-16 Hz) averaged between −1 to +1 s around the SWR for each IC. Each line represents the mean activity of all the electrodes averaged only on 10% of events with the highest weight for each IC (one-way ANOVA, *p<0.05). On the bottom, spatial distribution of the spindle power around −0.2 to 0.2s for the 10% group. **(C)** Spindle power data for each SWR were fitted with the IC weights as parameters in a GLM to observe the predominance of the spindle power in each electrode for the IC groups. **(D)** IC components from the Spindle power data around the SWRs (averaged −0.2 to 0.2s). The bars are the correlation r values between the weights from the sIC (ECoG) and the IC (VSD) (*p<0.05).

Taking the 10% of the events with highest weight for each IC, we observed that the overall spindle power is significantly higher for IC1 compared to IC2 and IC3 (Figure 4 B; one-way ANOVA, p-values: 10% = 0.04; One-way t-test compared to zero, p-value: 10% = IC1/0.001, IC2/0.61 and IC3/0.92). Taking 15% and 20%, which include more events that are poorer representatives of each IC, we observed a decay in the effect observed in IC1 (Figure S3 E; 15%: IC1 p = 0.003, IC2 p = 0.38 and IC3 p = 0.78. 20%: IC1 p = 0.004, IC2 p = 0.14 and IC3 p = 0.75).

More specifically, we found that spindle activity around IC1 also coincided with the topographic profile of the related IC (Figure 4 B; Figure S3 A).

In order to provide a concrete connection between the IC profiles and spindle activity, we also fitted a Generalized Linear Model (GLM) to observe the spatial patterns of spindles associated to each IC (Figure 4 C). For this, we used as predictors the scores of the IC’s. In contrast to IC2 and IC3 which showed almost no significant and low beta weights, again IC1 presented significant beta weights for the electrodes that occupy the activated regions in the IC profile (mostly RS). To specify this result as a property of the spindle frequency band, we also repeated the analysis for cortical Slow Gamma (20-45 Hz) and Medium Gamma (60-90 Hz; Figure S3 B). As opposed to spindle activity, neither Gammas showed significant activity around the SWR, nor clear spatial organization (Figure S3 C).

Next, we used the same ICA approach applied to the VSD data on the spindle power from the ECoG (Figure 4 D; Figure S3 D). Interestingly, the spindle independent components (sIC) showed a similar profile to the ICA components found in the VSD data. Correlating the scores from the sIC components with the cortical ICA components, we observed a positive correlation between the two ICs. However, this correlation is not significant.

### Cortical spindles in Retrosplenial cortex correlates with CA1 Slow Gamma activity during the SWR

Previously, we have shown that RS activity is connected to Slow Gamma (20-45 Hz) oscillations in the CA1 during NREM sleep (Pedrosa et al., 2022a). Furthermore, this interaction could also be observed with Slow Gamma bouts simultaneous with SWRs. This led us to ask whether cortical spindles also participate in this Retrosplenial-hippocampal interaction during Ripples. To observe how this Slow Gamma interacts with the cortical spindles during SWRs, we computed the cross correlation between the Slow Gamma power during the Ripple for each IC with the spindle power in each ECoG channel around the Ripple (Figure 5 A). For this correlation we used the 10% events with highest weight for each IC. We observed a significant correlation in the channel located at RS in IC1. Interestingly, the correlation between RS spindles with CA1 Slow Gamma peaked before the Ripple appearance. We did not find a significant result for IC3 and only a marginally significant site for IC2. This effect could not be observed when we correlated Ripple power with Spindle power across the ICs (Figure S4).

**Figure 5:**
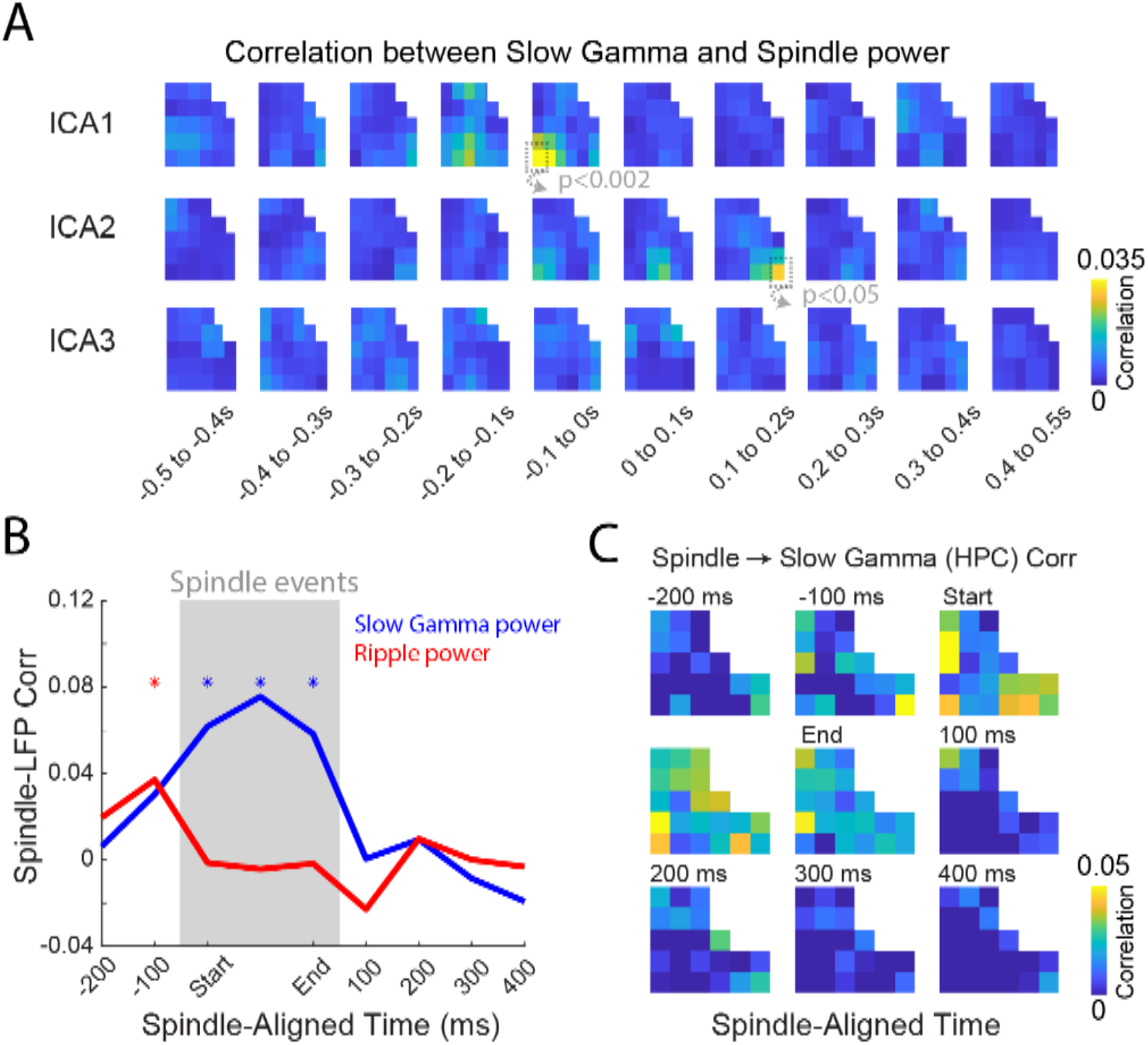
Slow Gamma during SWR is correlated to the spindle power in Retrosplenial cortex. **(A)** R values of the cross correlation between hippocampal Slow Gamma power with the spindle power in each channel around the Ripples events. The cross correlation was computed with the averaged spindle power in a 100 ms window around the Ripples. **(B)** Temporal correlations between the overall amplitude of cortical spindle events and Ripple and Slow Gamma power in the hippocampus, partialized by the power in the spindle frequency band in the hippocampus (*p<0.05). **(C)** Same as (B) but considering the correlation of hippocampal activity in the ripple or gamma bands with spindle activity in single ECoG channels.

We then took the opposite perspective and analyzed this cortico-hippocampal interaction at the time of spindle events. For this, we detected cortical spindles and computed the correlation between hippocampal Slow Gamma and Ripple power with the size of cortical spindle events. To minimize the influence of the volume conduction from the spindles, we partialized the correlation by the spindle frequency power (8-16 Hz) in the hippocampus. Interestingly, we found that the correlation between cortical spindles and Slow Gamma in the hippocampus showed significance during the entire duration of the spindle bout (Figure 5 B, p<0.01). On the other hand, Ripples only presented a small significant correlation on the period pre spindle. In particular, our results indicate that the correlation between cortical spindles and hippocampal Slow Gamma are spatially dominant on the cortical electrodes that overlap with the cortical DMN areas (Figure 5 C; Figure S4 B).

To investigate the CA1 source of this Gamma activity, we used a second dataset combining wide-field voltage sensitive imaging (VSI) with a high-density silicon probe in CA1 in mice expressing genetically encoded voltage indicators (GEVI) (Song et al., 2018) during natural NREM sleep (Figure 6 A-B), and repeated the same analysis of Figure 3. In this second dataset, we also showed that the Slow Gamma observed in the hippocampus is linked to the RS component and is mainly observed in the *stratum radiatum* layer (n=5 mice, Figure 6 C-E).

**Figure 6:**
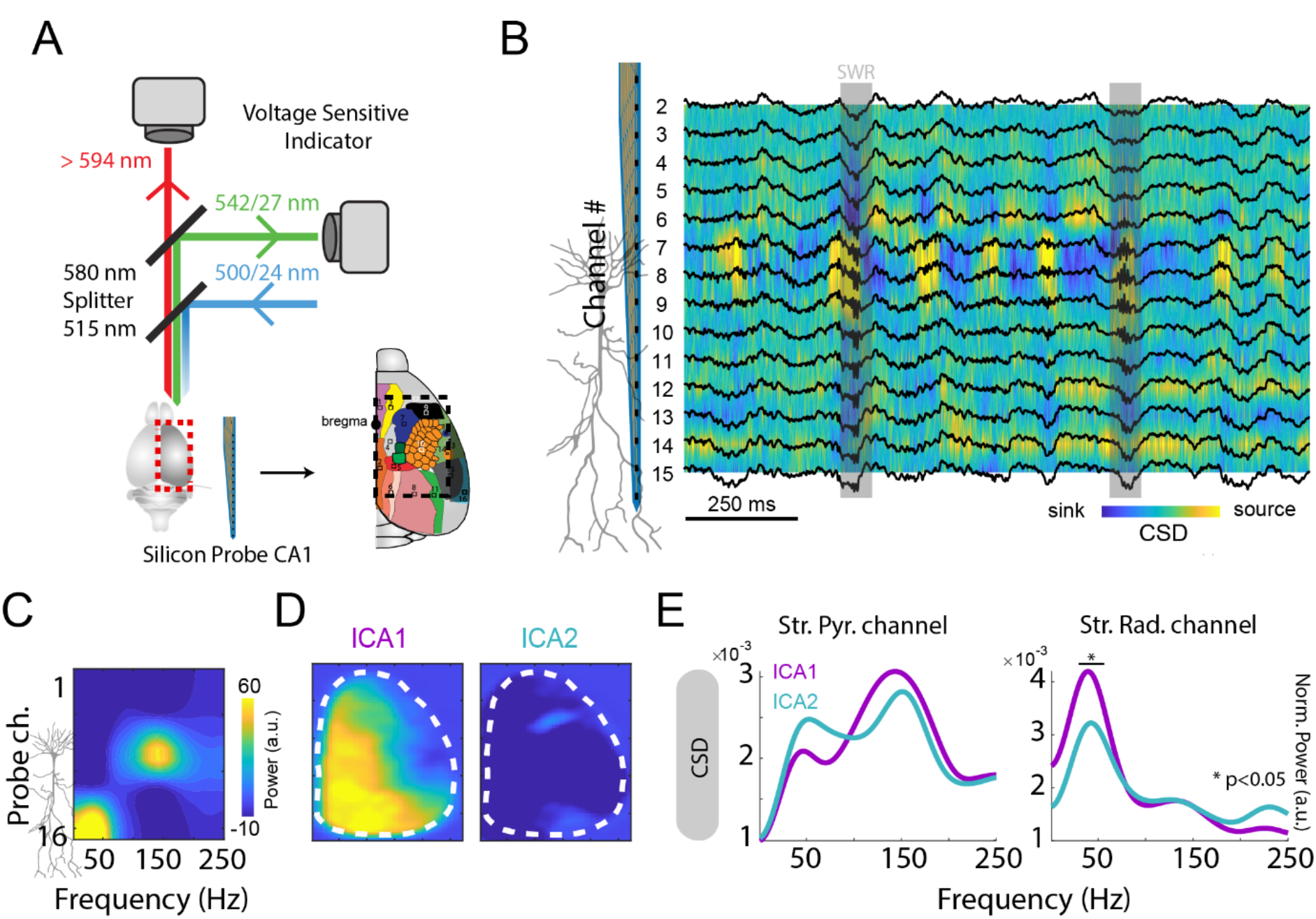
Hippocampal slow gamma power is most strongly correlated with DMN activity during SWR. **(A)** Illustration of the experimental setup used for the natural sleep recordings. In addition to the wide-field voltage imaging, we also implanted a high-density silicon probe (16 channels) in the hippocampus to record layer specific information in GEVI mice. **(B)** Current-source density (CSD) signal computed from the silicon probe data in different CA1 layers during NREM sleep. **(C)** Normalized power spectral activity across different CA1 layers for all the detected Ripples. **(D)** Cortical IC profiles calculated for the natural sleep recordings. Note we do not image A1/TeA in this dataset. **(E)** Frequency amplitude in Current Source Density (CSD) from channels located in *Str. Pyramidale (pyr.*) and *Str. Radiatum (rad.*). Note that no statistical difference was observed in Ripple frequency band in *Str. Pyramidale*, while in Slow Gamma the IC1 was significantly stronger in *Str. Radiatum* (one-sample t-test *<p<0.05, n= 5).

## Discussion

### The Default Mode Network emerges with different arrangements during hippocampal SWR

A basic tenet of the standard memory consolidation model is that the hippocampus encodes information and translates it to the neocortex (Marr et al., 1991; McClelland et al., 1995). Hippocampal SWRs are seen as critical for conducting this information to the neocortex and coordinating memory replays (Ji and Wilson, 2007; Peyrache et al., 2009). Hippocampal SWRs and UP states tend to co-occur in the neocortex (Battaglia et al., 2004), in particular in DMN areas (Kaplan et al., 2016; Karimi Abadchi et al., 2019; Pedrosa et al., 2022a). Parallelly, spontaneous cortical spindles transients (also located in the DMN) tends to precede SWR appearance (Siapas and Wilson, 1998). But besides them being statistically synchronous, the precise spatio-temporal patterns of coordination of SO, spindles and SWR have important implication for memory mechanisms (Geva-Sagiv and Nir, 2019; Skelin et al., 2021).

The topography of SO and spindles is not yet known with high resolution, partly due to the resolution limits. Here, we combined wide-field voltage imaging and ECoG from the neocortex with the CA1 LFP activity in mice during natural sleep and under urethane anesthesia (known as a good model of sleep) (Barthó et al., 2014; Clement et al., 2008; Greenberg et al., 2018; Pedrosa et al., 2022a). Voltage optical imaging offers significant advantages over classical approaches: (1) Temporal resolution comparable with electrophysiology and (2) no volume conduction.

Corroborating previous studies, our results also highlight the DMN as the most modulated network of the mouse brain around hippocampal SWRs (Kaplan et al., 2016; Karimi Abadchi et al., 2020; Pedrosa et al., 2022a). However, SWR events can coactivate with different subset of DMN areas (Figure 3). Analyzing the independent components, we observed that the fraction of the DMN concentrate around two main hubs: medial (RS, AC, Parietal and secondary Visual cortex) and lateral (A1, A2 and TeA). Previously, (Lu et al., 2012) identified these two distinct subsystems inside of the DMN, describing the medial network (centered around RS) as a more hippocampal-related module and the lateral network as a “viscerosensory” module due to the connection to multiple sensory modalities. In terms of functionality, these differences in activations across the DMN areas during SWRs may correspond to different information reactivated (Boyatzis et al., 2014). Thus, we may suggest that DMN activity configurations linked to different SWRs might be associated to the consolidation process of memories of distinct kinds.

### Hippocampal SWRs correlate to the RS “up-state” and spindle

RS cortex is one of the most strongly activated areas during hippocampal SWRs in sleep (Karimi Abadchi et al., 2020; Pedrosa et al., 2022a). Comparing the SWR features with the independent components from the neocortex, we found that exclusively IC1 (centered in RS) weights significantly correlates with the SWR duration. In fact, it has been shown SWR size is associated to cortical spindles during sleep (Ngo et al., 2020). IC1, also strongly involving RS, is the most correlated IC to SWRs (Figure 4). On the other hand, IC3 showed an inhibitory pattern across multiple cortical areas. This behavior has been previously observed during awake SWRs (Abadchi et al., 2022). Here we show that a fraction of sleep SWR are also related to cortical suppression, which includes spindle activity.

### Thalamocortical spindles in RS around hippocampal SWR

Within the imaged regions in dorsal cortex, RS emerged as the cortical area with the highest spindle amplitude during the SWRs. Sleep spindles are produced by a corticothalamic feedback (Kim et al., 2015; Pinault, 2004; Timofeev and Steriade, 1996). Here, we hypothesize that RS cortex may play a prominent role in the interactions between this thalamocortical and the hippocampus.

In humans, sleep spindles in the posterior cortex are commonly referred to as “centroparietal spindles”, due to the fact that they are observed via EEG channels on the surface of the parietal lobe (Andrillon et al., 2011; Cox et al., 2017; Gorgoni et al., 2016). Nevertheless, as the primary source of brain activity expressed in EEG is cortical regions close to the scalp and its ability to measure brain activity originating in deeper brain regions is limited, therefore finding the cortical origin of the spindle is challenging. However, activity from the RS (consisting of Brodmann areas 29 and 30, anatomically located deep in the parietal lobe; (Chrastil, 2018; Vann et al., 2009) would topographically map onto the centroparietal region of the brain where human studies often observe spindles, suggesting that RS may be crucial for the spindles measured in the centroparietal region of the cortical surface.

### Temporal relationship between RS and hippocampal activity around SWRs

Spontaneous neocortical activity during sleep statistically tends to organize in three functional networks (DMN, lateral network and somatomotor network; (Pedrosa et al., 2022a)). Within the DMN cortical areas, RS cortex is a focus of activity initiation (Karimi Abadchi et al., 2020). Importantly, transient activations starting from RS cortex are associated with a greater increase in Slow Gamma power in the hippocampus, accounting to a large extent for the correlations between SWR and cortical activity. Additionally, using a pseudo-causality analysis, <u>Pedrosa et al., 2022</u> also shows that in this RS-Slow Gamma (hippocampal) interaction the neocortex tends to precede the hippocampus. In contrast, ripples precede widespread neocortical “up-state” across multiple areas (Nitzan et al., 2020; Pedrosa et al., 2022a; Tong et al., 2021). These timing relationships, together (RS to Slow Gamma to Ripple to cortex) describe a circular mechanism evolving cortex and hippocampus around SWRs that may be fundamental for memory consolidation (Helfrich et al., 2019; Rothschild et al., 2017). To observe how cortical spindles may participate in this cortico-hippocampal schema, we first measured the hippocampal Slow Gamma power component correlation with the loadings of each ICs (Figure 5 A). Interestingly, IC1, centered in RS presented a statistically significant correlation. No correlation was observed between spindles and hippocampal ripple power (Figure S4).

In sum, our work presented a novel picture on the cortico-hippocampal relationship during SWR’s, indicating that RS cortex “up-state” and spindles activities potentially moderate a sequence of events during the hippocampal SWRs (Figure 7).

**Figure 7:**
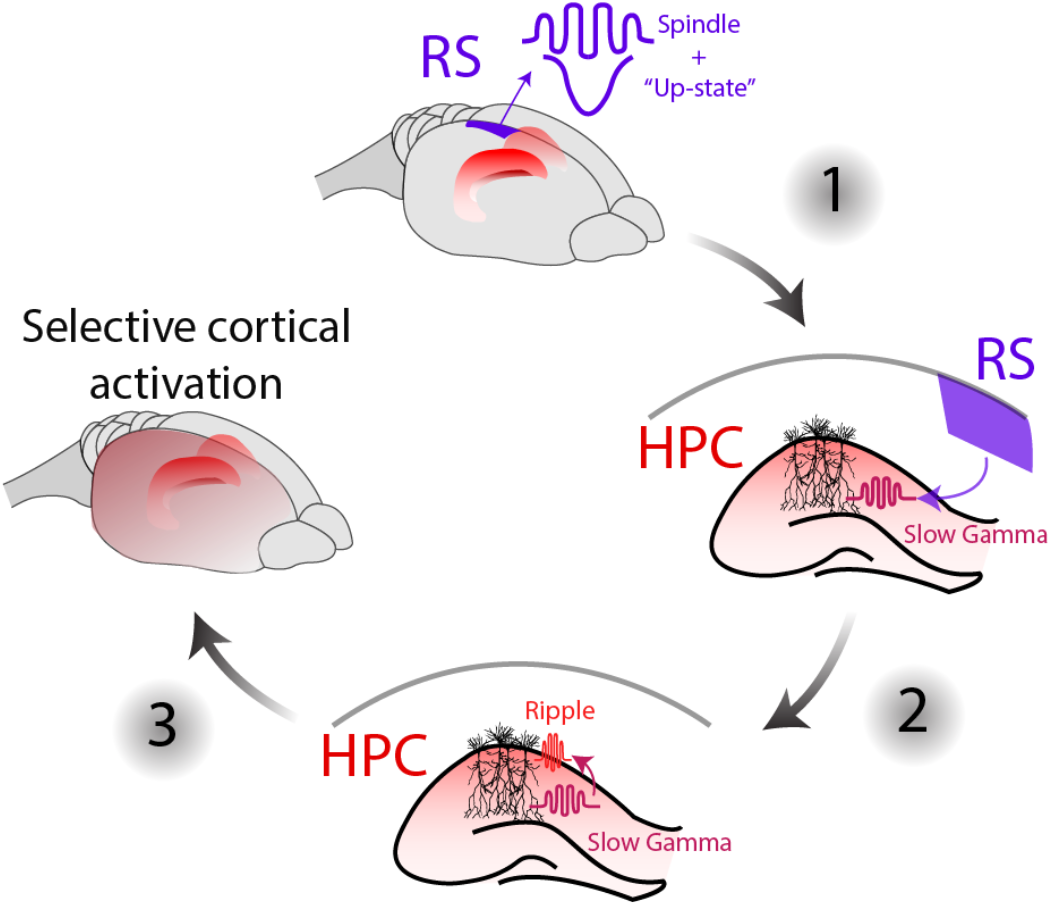
Schematic of the hypothesized sequence of events evolving the cortico-hippocampal interaction around the SWR.

## Materials and methods

### Animals

For acute VSD imaging (urethane group), five C57/Bl6 adult male mice (ages 2 to 4 months) were used. Under a 12-hour light/dark cycle, the animals were housed in conventional plastic boxes with other animals. Water and food were available to the animals at all times, ad libitum. The Animal Welfare Committee of the University of Lethbridge accepted the animal experimental methods and followed all of the standards set out by the Canadian Council on Animal Care (CCAC). They were given a pulse of electrical, tune or light stimulation to map different sensory cortices.

For VSI imaging, five CaMK2A-tTA;tetO-chiVSFP mice (male, 3 to 6 months old) were used, weighing between 25 and 35 g. To protect the implants, the animals were housed in groups until surgery day, then separately afterward. For this group, all the experiments were conducted during the light period. The Central Commission Dierproeven (CCD) and the Radboud University Animal Welfare Board approved this experimental technique, which was carried out in compliance with the Experiments on Animals Act and the European Directive 2010/63/EU on animal research.

### Surgery

#### Surgical preparation for the VSD imaging

First, mice were anesthetized with 3% isoflurane with 100% of oxygen and sedated with urethane (1.25 mg/kg). The animals were subsequently placed in a stereotaxic and kept under 0.5-1% isoflurane anesthesia using a nasal mask. A heating pad controlled by a feedback thermistor kept the body temperature at 37 °C. To reduce discomfort, 80 mg of dexamethasone was given intramuscularly. Following previous protocols (Karimi Abadchi et al., 2020; Pedrosa et al., 2022b, 2022a), two Teflon-coated stainless-steel wires (277 m diameter) were inserted as ground on the pial surface of the cerebellum and in the hippocampus (−4.3 mm AP and +2.4 mm ML, +1.8 mm DV at a 60-degree angle) for hippocampal registration. Super glue and acrylic resin were used to secure the electrodes to the skull. Next, a tracheostomy was performed to aid the animal’s breathing. For skull removal, a unilateral craniotomy window was created. Following the craniotomy, the exposed cortex was fully covered with 4-(2-hydroxyethyl)-1-piperazineethanesulfonic acid (HEPES) - buffered saline solution (1 mg ml1) and administered to preserve hydration.

#### Surgical preparation for the VSI group

Animals were sedated with 2% isoflurane at 100% oxygen concentration, then placed in a stereotaxic apparatus and kept under 0.5-1.5 percent isoflurane via a nasal mask. A heating pad controlled by a feedback thermistor kept the body temperature at 37 °C. At the start of the anesthetic, a subcutaneous dose of 5 mg/kg of carprofen was injected, followed by 50 μl of 2 percent lidocaine through the scalp. During the surgery, the levels of isoflurane and oxygen were monitored and regulated to keep the respiratory rate between 0.5 and 1.5 Hz. A skull screw was then placed above the left cerebellum to provide a shared reference and ground for the silicon probe. After that, two more skull screws were placed on the left hemisphere to help secure the head-plate implant. The exposed right hemisphere skull was next thinned to remove tiny skull capillaries in order to improve later voltage imaging recording. We performed a craniotomy on the pial surface of the right cerebellum (−4.3 mm AP and +2.4 mm ML) and inserted a 16 channel silicon probe with an inter-site pitch of 50 m (E16 R 50 S1 L10, Atlas Neuroengineering, Belgium) in the intermediate hippocampus (+2.1 mm DV at a 57-degree angle towards the back) for CA1 laminar recordings. The electrodes were carefully attached to the skull with cyanoacrylate. Finally, we used acrylic cement to attach a head-plate with a wide field view of the right hemisphere to the skull (Super-Bond C&B). All animals were given at least four days after surgery to begin acclimating to the arrangement. For a complete description of the protocol, see (Pedrosa et al., 2022b).

### VSD imaging recordings

After the surgical procedure, RH1961 was dissolved in 0.5 mg/ml HEPES-buffered saline and administered to the exposed cortex for 40-60 minutes. After that, 1.5% agarose in HEPES-buffered saline was applied over the cortex and sealed with a coverslip to minimize respiration artifacts. Using a CCD camera (1M60 Pantera, Dalsa, Waterloo, ON) and an EPIX E8 frame grabber controlled by XCAP 3.7 imaging software (EPIX, Inc, Buffalo Grove, IL) and XCAP 3.8 imaging software (EPIX, Inc, Buffalo Grove, IL), VSD image were acquired in 12-bit format at a frame rate of 200 Hz (EPIX, Inc.). Images were captured using a macroscope with front-to-front video lenses (8.6 x8.6 mm field of view, 67 μm per pixel). Two LEDs (627-nm center, Luxeon K2) were used to excite the VSD, and the fluorescence emitted was filtered through a 673-nm to 703-29 nm bandpass emission filter. To characterize the neocortical topography, we applied sensory stimulation to localize different primary sensory areas (HLS1: hidelimb, FLS1: forelimb, V1: visual cortex) in order to inspect the local evoked activity in the VSD data (Mohajerani et al., 2013).

### VSI recordings (natural sleep)

Two synchronized sCMOS cameras (PCO edge 4.2) operated by a TTL external trigger and coupled with a Leica PlanAPO1.6 lens were used to capture widefield epifluorescence. The excitation light was provided by a high-power halogen lamp (Moritex, Brain Vision) installed in the macroscope. As optical filters we used: mCitrine emission FF01-542/27, mCitrine excitation 500/24, mKate2 emission BLP01-594R-25 and beam splitters (515LP,580LP). The image was captured at 375×213 pixels (spatial resolution of ~36 um/pixel; 12 bit) at 50 Hz in 15 minutes blocks.

### ECoGs

The fabrication of the PEDOT:PSS ECOGs (Figure S1 D) has been described previously (Donahue et al., 2018; Khodagholy et al., 2013) and consisted in the deposition and the patterning of parylene, metal and PEDOT:PSS. First, a 3 μm-thick parylene-C film was deposited on top of a glass slide using a SCS-Labcoater 2. Metal pads and interconnects were patterned by lift-off using S1813 photoresist exposed to UV-light with a SUSS MBJ4 mask aligner and developed with MF-26 developer. Chromium (5 nm) and gold (150 nm) were deposited with a thermal evaporator and the final pattern was obtained by lifting the resist in a mixture of acetone and isopropanol. A second 3 μm-thick layer of parylene was used to insulate the metal tracks, followed by the deposition of a soap solution (4%wt) and a third “sacrificial” 3 μm-thick layer of parylene. They were patterned using AZ9260 photoresist and reactive ion etching with an oxygen plasma using an Oxford 80 plus, leading to openings on the electrodes (20 μm x 20 μm) and the contact pads.20 mL of PEDOT:PSS (Clevios PH 1000) were mixed with ethylene glycol (5 mL) and dodecyl benzene sulphonic acid (50 μL) to improve the conductivity and 3-glycidoxypropyltrimethoxysilane (0.25 g) to prevent film delamination. The solution was spin-coated on the glass slides and the sacrificial layer of parylene was removed by peel-off to leave PEDOT:PSS only on contact pads and electrodes openings. Finally, the samples were baked at 140 °C for 1 hour to anneal the PEDOT: PSS films then immersed in water to remove excess of PSS in the PEDOT:PSS and release the ECOGs from the glass slides.

### CA1 LFP and ECoG recordings

We employed a transparent ECoG grid to obtain an electrophysiological profile of the neocortex. The recording sites on the grid are grouped in a 6×5 matrix (each site is spaced 1 mm apart) and cover a total area of 6 mm x 5 mm. To capture neocortical activity, we first grounded the grid to the animal’s nose with a subcutaneously implanted electrode. The ECoG matrix was then placed across the mouse’s right neocortex, encompassing the same wide-field as the optical imaging. After that, we applied HEPES-buffered saline to the exposed cortical region to keep it hydrated. The electrophysiological data were captured using 32-channel headstages (Intan Technologies’ RHD2132) coupled to an Open Ephys recording system. The raw signals were recorded at 20 kHz sampling rate and filtered between 0.1 and 7500 Hz with a pre-amplifcation of 20x.

For the VSD recordings, we recorded the hippocampus LFP at 20 kHz sampling rate in a Digidata 1440 (Molecular Device Inc, CA) data recording system. The recorded data were filtered (0.1-10000 Hz) using a Grass A.C. Model P511 (Artisan Technology Group, IL). While for the VSI recordings we used a 16 channels silicon probe (Atlas Neuroengineering, Belgium). The CA1 profile data were acquired using a headstages (RHD2132, Intan Technologies) with an Open Ephys system. The signals were recorded at 20 kHz sampling rate and filtered between 0.1 and 7500 Hz.

### Voltage imaging preprocessing

For the VSD imaging preprocess we used the steps from previous work (Karimi Abadchi et al., 2020; Nazari et al., 2022). We first applied a high-pass Chebyshev filter above 0.2 Hz to the time series of each individual pixel. Next, we averaged the filtered signal of each pixel and defined it as the baseline signal (F_0_). The filtered signal was then subtracted by the F_0_, and the resultant difference was divided by the Fo values (F-F_o_/F_0_ × 100). For the VSI recording and preprocessing details we repeated the protocol previously reported (Pedrosa et al., 2022b, 2022a; Song et al., 2018).

### Ripple detection

We initially downsampled the data to 1 kHz and filtered in between 150-300 Hz. After, we detected the peaks of the envelope filtered data that crossed a threshold between 3 and 5 s.d. plus the mean. Based on the points that crossed 1.5 s.d. before and after the detected points, the start and end of the ripple were determined. Intervals above 30 ms and shorter than 300 ms were considered ripple events. In addition, the time between the end of one ripple and the beginning of the next had to be at least 100 ms.

### Spindle detection

We filtered the raw signal from each ECoG channel fin the 8-16 Hz band, computed the power of the resulting signal and z-scored it. For each channel we defined the presence of significant spindle power whenever this score exceeded the value of 2. We then defined a measure of global spindle power activity as the number of simultaneously ‘active’ ECoG channels at any moment in time. Finally, we classified spindle events as time windows in which the number of active channels exceeded the average value computed on the entire recording session. We imposed a minimum duration to accept any of these events at 200ms. For each of these events we then computed a size score as the cumulative sum over the duration of the event, of the number of active channels per time bin. A similar measure was also computed for each channel separately, as the number of time bins within an event in which that particular channel surpasses the spindle detection threshold.

We then correlated the appearance of spindle events in the cortex with the presence of activity in specific frequency bands in the hippocampus. Hippocampal LFP was filtered in separate frequency bands: spindle (8-16 Hz); gamma (25-50 Hz); ripple (150-250 Hz). We then computed the power in time for each of these bands. To obtain an estimate of the ripple power density, the ripple time series was also smoothed with a large temporal kernel of 100ms.

The temporal alignment between cortical and hippocampal oscillatory events was evaluated by normalizing the length of all spindle events to a time window of equal length and subsequently computing the correlation between the size of each aligned spindle event and the hippocampal power detected at different lags with respect to the onset and offset of the spindle event. In particular we computed the correlation of gamma power vs. spindle size and ripple power vs. spindle size. In both cases we factored out the contribution of the spindle band in the hippocampus (a potential confounding factor due to volume conductance between cortical recording sites and the hippocampus) by computing a partial correlation of the aforementioned factors. Such correlations were computed both considering the overall size of the cortical spindle event (thus including all ECoG channels) or separately by considering the presence of spindle-related activation in each of the channels separately.

### Spindle power

Spindle power was defined as the envelope of the filtered Raw data (8-16 Hz) in each individual channel. In order to avoid problems with the different impedance across electrodes, the resultant signal was Z-scored for each individual channel.

### Register Multimodal Images

To map data from all mice to a common spatial framework, the *imregister* function in the Image Processing toolbox (https://nl.mathworks.com/products/image.html) was applied. This function uses an intensity-based image registration to automatically align images to a common coordinate. For this, we re-spaced all the images recorded for all the animals to a common mask.

## Author contributions statement

Projected conceived and designed by R.P., M.N., M.M and F.B.; R.P. and F.S. analyzed the data; R.P. and M.N. performed surgery and electrophysiological recordings; L.K. and C.B. designed and manufactured ECoG grids; R.P. prepared figures; R.P. and F.B., draft manuscript; R.P, M.M., L.K., C.B, F.S and F.B., edited and revised manuscript.

## Acknowledgments

This work was supported by Natural Sciences and Engineering Research Council of Canada 819 (grant no 40352 & 1631465 to M.H.M.), Alberta Innovates (M.H.M.), 820 Alberta Prion Research Institute (grant no. 43568 to M.H.M.), and Canadian Institute for Health Research (grant no 390930 & 156040 to M.H.M.), National Science Foundation (M.H.M.) and European Commission grants grants ERC-AdG 833964 “REPLAY_DMN” (to F.P.B.), MSCA ITN 765549 “M-GATE” (to F.P.B.), MSCA Intraeuropean Fellowship 840704 “BrownianReactivation” (to F.S. and F.P.B.)

The authors thanks Thomas Knöpfel for donating the Butterfly 1.2 mice and for advice on imaging systems and preprocessing. The authors also thank Dr. Jianjun Sun at the University of Lethbridge for the surgical assistance.

## Conflict of interest statement

The authors declare no conflict of interest in this paper.

## Supplementary Material

**Figure S1:**
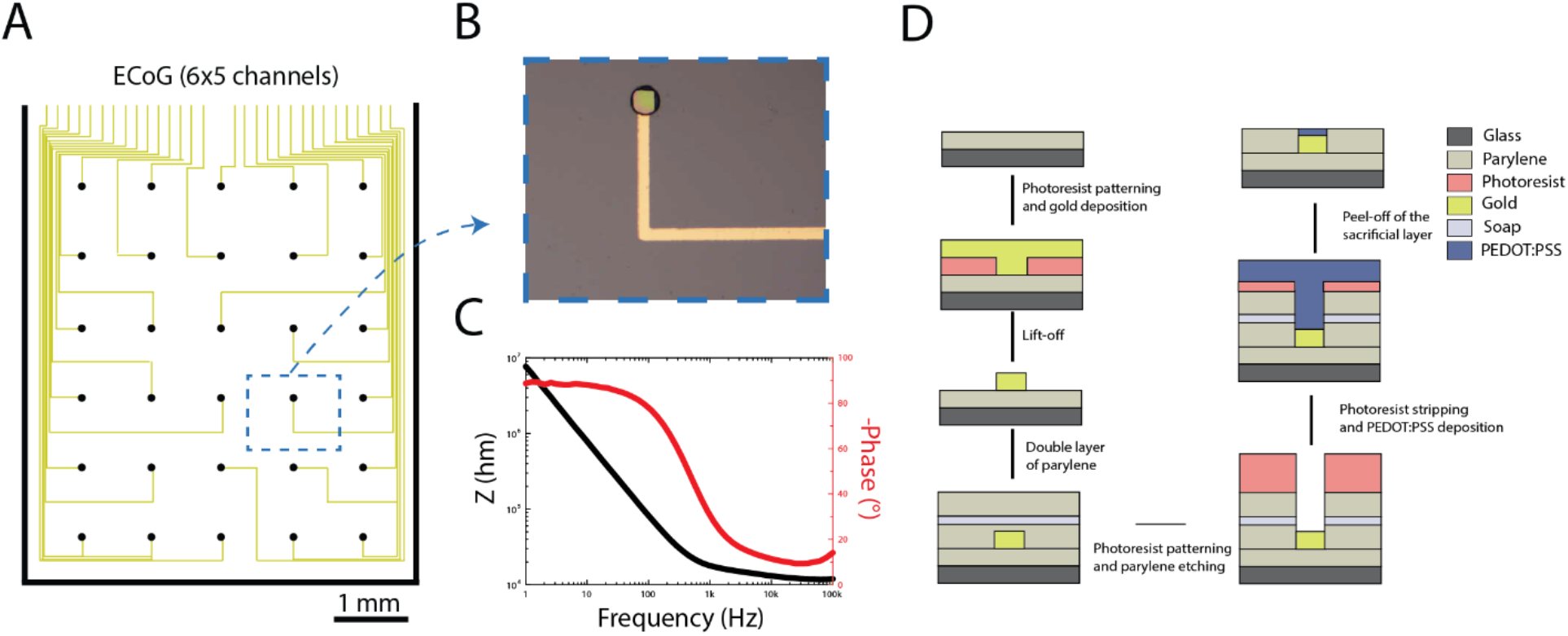
**(A)** Schema of the 30-channel ECOGs. **(B)** Magnified picture of 20 μm^2^ x 20 μm^2^ electrode covered with PEDOT:PSS. **(C)** Electrochemical impedance magnitude and phase of a single electrode. At 1 kHz, the impedance is about 20 kΩ. **(D)** Schematic of the microfabrication process of the ECOGs.

**Figure S2:**
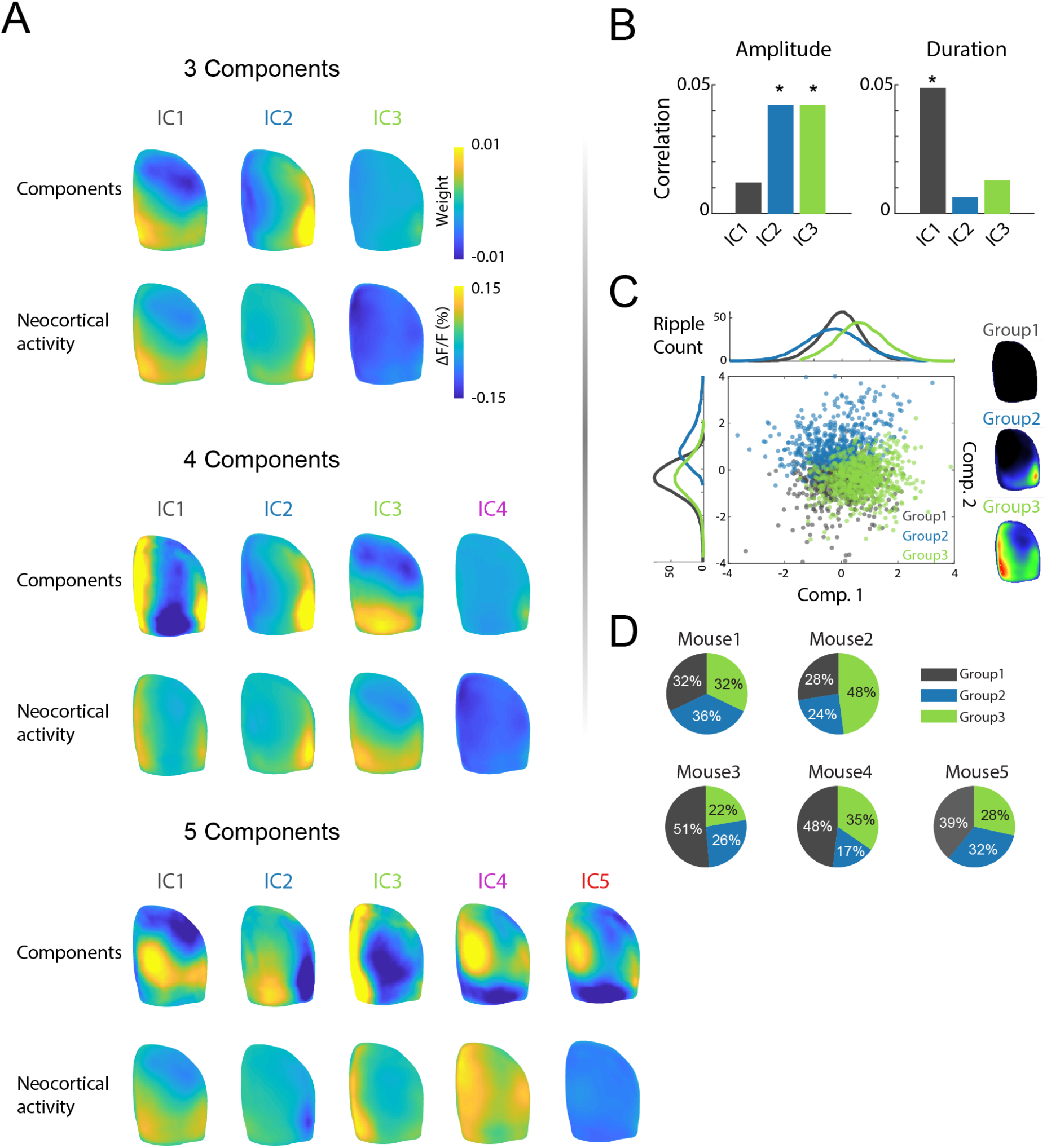
Neocortical activity around Ripples (−0.2 to 0.2s) classified in 3, 4 or 5 IC components. **(A)** The top plots are the components of the IC, while the bottom shows the average neocortical activity of the 20% of the total Ripple events with the highest weight value for each component. **(B)** SWRs information for each IC. The bars show the correlation between the weight in each IC with the amplitude and duration of the Ripple events (* p<0.05). **(C)** Scatter plot shows the first two estimated independent components of the cortical activity for each detected SWR. In gray, green and blue, we show 3 groups classified by k-means using the first 3 estimated independent component. On the right, average of the cortical activity of each group. **(D)** Probabilistic distribution of the 3 groups for each individual mouse.

**Figure S3:**
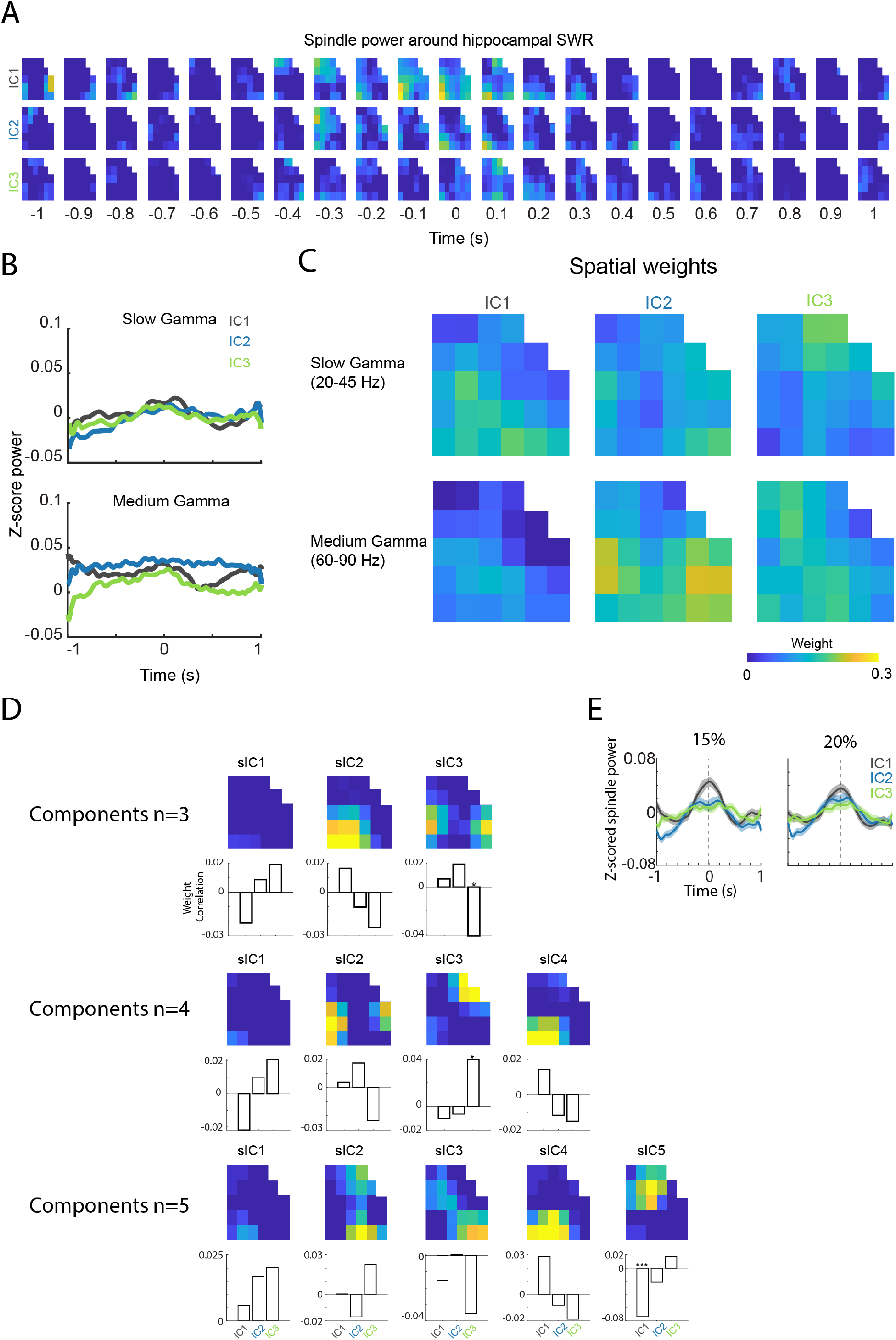
**(A)** Spatiotemporal Spindle power for each IC. The averaged events are the events with the highest weight for each IC (20% highest events) **(B)** Global Slow (20-45 Hz) and Medium (60-90 Hz) Gamma powers averaged between −1 to +1 s around the SWR for each IC. Each line represents the mean activity of all the electrodes averaged by 20% of the events with highest weight for each IC. **(C)** GLM of Slow and Medium Gamma power data. The parameters used here are the weights of the IC’s. **(D)**. IC of the Spindle power around the Ripples (−0.2 to 0.2s). On the top we show each component. On the bottom, the bars represent the correlation between the weight from the neocortical IC components with each spindle IC component (sIC). We computed the analysis for 3, 4 and 5 number of components (p*>0.05 and p***>0.0005). **(E)** The global spindle power (8-16 Hz) averaged between −1 to +1 s around the SWR for each IC. Each line represents the mean activity of all the electrodes averaged by the respective 15% and 20% events with highest weight for each IC (one-way ANOVA, both results are non-significant).

**Figure S4:**
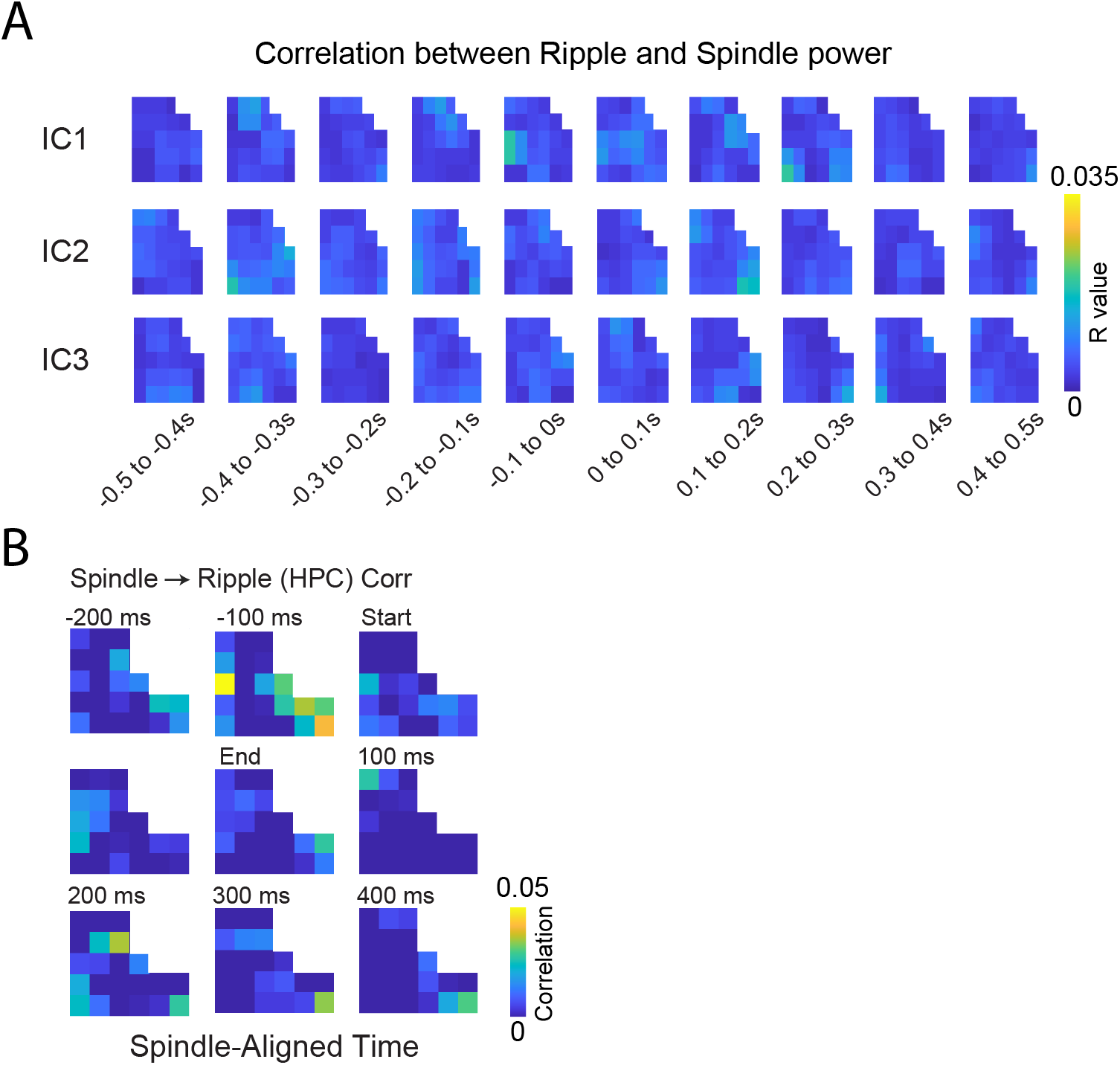
**(A)** R values of the correlation between Ripple power during the Ripple events with the spindle power in each channel around the Ripples. The correlation for each IC was computed in the 20% of the events with highest weight for each IC. **(B)** Spatiotemporal correlation between cortical spindle and Ripple in the hippocampus partialized by the spindle frequency band in the hippocampus.

## Notes

### Competing Interest Statement

The authors have declared no competing interest.

